# SSVEPs reveal dynamic (re-)allocation of spatial attention during maintenance and utilization of visual working memory

**DOI:** 10.1101/2023.08.29.555110

**Authors:** Samson Chota, Arnaud Bruat, Stefan Van der Stigchel, Christoph Strauch

## Abstract

Visual Working Memory (VWM) allows us to temporarily store goal-relevant information to guide future behavior. Prior work has established that VWM and spatial attention are intrinsically connected – even if location is irrelevant for responses: behavioral and neural correlates of attention show spatial biases, specific to the location at which memory items were encoded. This suggests that VWM is spatially organized and that maintaining information might rely on the allocation of spatial attention towards the location of memory items. Importantly, attention often needs to be dynamically redistributed between several locations, e.g. in preparation for an upcoming probe. Very little is known about how attentional resources are distributed between multiple locations during a VWM task and even less about the dynamic changes governing such attentional shifts over time. This is largely due to the inability to use behavioral outcomes to reveal fast dynamic changes within trials.

We here demonstrate EEG Steady-State Visual Evoked Potentials (SSVEPs) to successfully track the dynamic allocation of spatial attention during a VWM task. Participants were presented with to-be-memorized gratings and distractors at two distinct locations. During maintenance and retrieval, each location was tagged with a disk flickering at either 10 or 13 Hz. This allowed us to dynamically track attention allocated to memory and distractor items via their coupling with space by quantifying the amplitude and coherence of SSVEP responses in the EEG signal to flickering stimuli at the former memory and distractor locations.

SSVEP responses at memory locations did not differ from distractor locations during early parts of the maintenance window. However, shortly before probe comparison, we observed a decrease in SSVEP coherence over distractor locations indicative of a reallocation of spatial attentional resources. Reaction times were shorter when preceded by stronger decreases in SSVEP coherence at distractor locations, reflecting the reallocation of attention from the distractor location to the memory location or towards the distinct upcoming probe location.

*Broader Significance* We demonstrate that SSVEPs can inform about dynamic processes in VWM, even if location does not have to be reported by participants. This finding not only supports the notion of a spatially organized VWM, but also reveals that SSVEPs betray a dynamic prioritization process of working memory items and locations over time that is directly predictive of memory performance.

## Introduction

Visual Working Memory (VWM) allows to temporarily store information such as the appearance and location of objects that are presumed relevant for future behavior (Baddeley, 1992; Logie & Marchetti, 1991). While attention is an essential component of VWM, little is known about how its spatial allocation changes over time during maintenance and utilization of memoranda (Awh et al., 2006). Past work has highlighted that working memory is spatially grounded, by demonstrating that spatial attention is automatically biased towards memorized locations (Awh & Jonides, 2001). Spatial attention and working memory likely share neural mechanisms, as drawing spatial attention away from memorized locations results in impaired reports (Awh et al., 1998; Awh & Jonides, 2001; Smyth, 1996).

Intriguingly, memory location-specific effects on attention can also be observed *when the location itself is irrelevant* e.g., when only the color, but not the location, of an item needs to be reported (Theeuwes et al., 2011; but see also Awh et al., 1998). So far, these effects have been studied with so-called retro-cues, cues that direct attention to memoranda after templates have been taken away. Retrocued items are associated with better recall performance (Souza & Oberauer, 2016). In a recent series of experiments, participants were tasked with memorizing the orientation of two colored bars presented in the periphery, after which color retro-cues at fixation indicated which item will likely be subsequently probed (van Ede et al., 2019, 2020, 2021). Results showed that retro-cues led to gaze biases towards the location of cued items, even though that location did not have to be remembered nor did the memory probe ever appear in that location. Such directional biases in gaze can track the direction of covert spatial attention (Engbert & Kliegl, 2003; Kustov & Lee Robinson, 1996; Schall & Hanes, 1993; Zhou & Desimone, 2011) and are assumed to index attentional selection within working memory. Location is therefore likely an essential component of VWM representations with location taking a grounding role in VWM storage (Schneegans & Bays, 2017; van Ede et al., 2019). This conclusion is further supported by electrophysiological evidence demonstrating that markers of attentional selection such as alpha lateralization (Ede et al., 2017; Foster et al., 2017; Liu et al., 2022, 2023; Poch et al., 2014, 2017) and the N2Pc event-related potential (Dell’Acqua et al., 2010; Kuo et al., 2009) as well as markers of VWM storage such as contralateral delay activity (CDA, Eimer & Kiss, 2010) show similar spatial biases. We here extend this toolbox by utilizing SSVEPs as a continuous measure of spatial attention.

It is becoming increasingly evident that visual working memory needs to be viewed as a dynamic and flexible process reflecting the fast changing environment in which it is utilized (Buschman & Miller, 2022; Chota & Van der Stigchel, 2021; Nobre & van Ede, 2023). Novel methods capable of capturing fast changes in the prioritization of memoranda as well as their interaction with the external world are therefore urgently needed. We therefore first set out to **test whether SSVEPs allow to study for spatiotemporally dynamic internal operations in VWM** . The findings outlined above suggest that during VWM maintenance, spatial attention is distributed among memory items resulting in location-specific enhancements in perceptual processing. Several questions, however, remain unanswered. Second, **how are attentional resources *spatially* distributed between multiple relevant or irrelevant locations during a VWM task**This question has proven difficult to answer using classical methods of measuring covert attention such as microsaccades. For instance, it was recently demonstrated that microsaccades are directed towards the midpoint between two items, irrespective of one or both being cued as behaviorally relevant (Willett & Mayo, 2023), showing a limitation of spatial specificity of microsaccades in some cases. Furthermore, prior behavioral studies utilized dual-tasks during the maintenance period to probe spatial attention which could be a potentially confounding factor as these tasks themselves require allocation of spatial attention to memory locations, e.g. to detect a target (Awh & Jonides, 2001; Golomb et al., 2008). Third and foremost, what is the temporal dimension of this hypothesized spatial distribution? In other words, **how is spatial attention (re)allocated between memory, distractor, or probe locations *over time*?**

We used EEG Steady-State Visual Evoked Potentials (SSVEPs), rhythmic brain responses to flickering stimuli, that have been shown to increase in amplitude as a result of covert attention allocated to the flickering stimulus location (Gulbinaite et al., 2017, 2019). They allow for direct and highly time-resolved measures of spatial attention and ensuing facilitation of sensory processing, without the need for a secondary behavioral task. In the current experiment participants were presented with two tilted gratings at distinct spatial locations. Participants were instructed to memorize the identity but not location of both (memory + memory) or only one (memory + distractor) grating. We subsequently presented two flickering discs over both locations which allowed us to measure the magnitude of SSVEP responses (spatial attention) allocated to memory and distractor locations during maintenance and preparation for probe comparison. This enabled us to compare how attention was dynamically allocated between one or two locations previously occupied by memory items or distractors.

We showcase that SSVEP responses are a powerful new approach to quantify dynamic changes in internal attention towards individual items in working memory. We found enhanced SSVEP responses at memory compared to distractor locations, supporting the notion of a spatially organized and spatially selective VWM. Crucially, time-resolved analysis of SSVEP responses showed a decrease in attention directed towards distractor locations shortly before probe presentation, indicating a reallocation of attention towards the upcoming central probe location. This decrease did not occur for memory locations and predicted subsequent reaction times: participants with a stronger attentional reallocation/suppression of the distractor location responded faster to probes. This shows that the flexible use of limited attentional resources during VWM is key for efficient task performance. Furthermore, we found that SSVEP responses were not reduced when attention was distributed between two memory locations as compared to one, potentially because in the latter attentional resources were distributed equally between memory and distractor locations before being reallocated shortly before probe presentation. Our findings confirm accounts of a spatial organization of VWM and demonstrate that VWM maintenance and utilization is accompanied by dynamic, location-specific and behaviorally relevant (re-)distribution of spatial attention.

## Materials and Methods

### 1. Participants

N = 26 participants (14 female, 12 male) with normal or corrected to normal vision enrolled in the experiment. None of the participants reported a history of psychiatric diagnosis. Informed consent forms were signed before the experiment. The study was carried out in accordance with the protocol approved by the Ethics Committee of the Faculty of Social and Behavioural Sciences of Utrecht University and followed the Code of Ethics of the World Medical Association (Declaration of Helsinki). Participants were compensated with 10 €/hour.

### 1. 2. Stimuli

Memory and probe stimuli consisted of black and white oriented gratings (diameter: 4° dva, spatial frequency: 4 cpdva) whose orientation was randomly selected from six frequencies on every trial (12, 42, 72, 102, 132, 162°). Memory orientations in the two-item condition were always distinct. Distractors used in the one item condition were created using a Mondrian Mask (diameter: 4° dva, code from Christophel et al., 2012). A large rectangular mask using the identical Mondrian pattern was presented centrally following the memory and distractor items to prevent afterimages (height: 5° dva, width: 8° dva). SSVEP stimuli consisted of circular discs (diameter: 4° dva) and were sinusoidally modulated at a frequency of 10 Hz and 13.333 Hz. These frequencies were chosen to allow for a precise estimation of their power within a 1.5 s window.

### 1. 3. Protocol

Stimuli were presented on an LCD display (27-inch, 2560 x 1440 resolution, 120 Hz refresh rate) using the Psychophysics Toolbox (Brainard, 1997; Kleiner et al., 2007) running in MATLAB (MathWorks). Participants were seated at 58 cm from the screen on a chinrest to prevent excessive head movements.

Participants performed a delayed match-to-sample task, in which they reported whether the orientation of the probe matched either of the initially memorized stimuli. At the beginning of each block participants were informed about the number of items they were required to memorize in this particular block (“One Item Condition” versus “Two Item Condition”). The starting block of individual participants was randomized and item conditions alternated between blocks. Each block contained 44 trials and participants performed a total of 14 blocks (2 training blocks, 88 trials total + 12 experimental blocks, 528 trials total).

Trials began with a central black fixation cross presented for 1500 ms on gray background. A sequence of two items (one item condition: distractor + memory, two item condition: memory A + memory B) was presented to the left and right side of fixation in a randomized order (eccentricity: 3° dva, figure 1). The first item was presented for a random duration between 200 and 500 ms (steps of 30 ms), followed by the second item which was presented for a duration of 100 ms. The presentation time of the first item was varied to prevent participants from developing precise temporal expectations for the onset of the second stimulus. The temporal order and position of memory and distractor presentation was randomized. After a screen containing only the fixation cross (200 ms), a large rectangular mask (500 ms) was presented to prevent afterimage effects on the SSVEP responses. This mask was immediately followed by two circular SSVEP entrainers whose position overlapped with the original location of the memory items and/or distractors. The SSVEP stimuli flickered for a variable delay of 2,000 to 2,330 ms (in steps of 30 ms) with 13.333 Hz always being presented on the right and 10 Hz always being presented on the left side. SSVEP duration was varied to prevent participants from developing precise temporal expectations on the appearance of the probe. The location of the flicker frequencies was kept identical throughout the task to allow for the inclusion of all trials in the Rhythmic Entrainment Source Separation procedure (*RESS*), which significantly increases the signal to noise ratio for the construction of the spatial filters (see methods section *5. RESS*). Subsequently, the probe was centrally presented for 50 ms, either matching the orientation of one of the initially presented items (75%) or matching none of the items (25%). In the case of a non-match its orientation was randomly drawn from one of the remaining orientations not used for a sample. Participants were given 1,500 ms to report either a match (keyboard button “P”) or a miss-match (keyboard button “Q”) after which they received feedback. Feedback was presented in the form of the fixation cross turning green (hits/correct rejections) or red (misses/false alarms) for 100 ms. If participants did not respond within 1,500 ms a message prompted them to respond faster.

**Figure 1.**
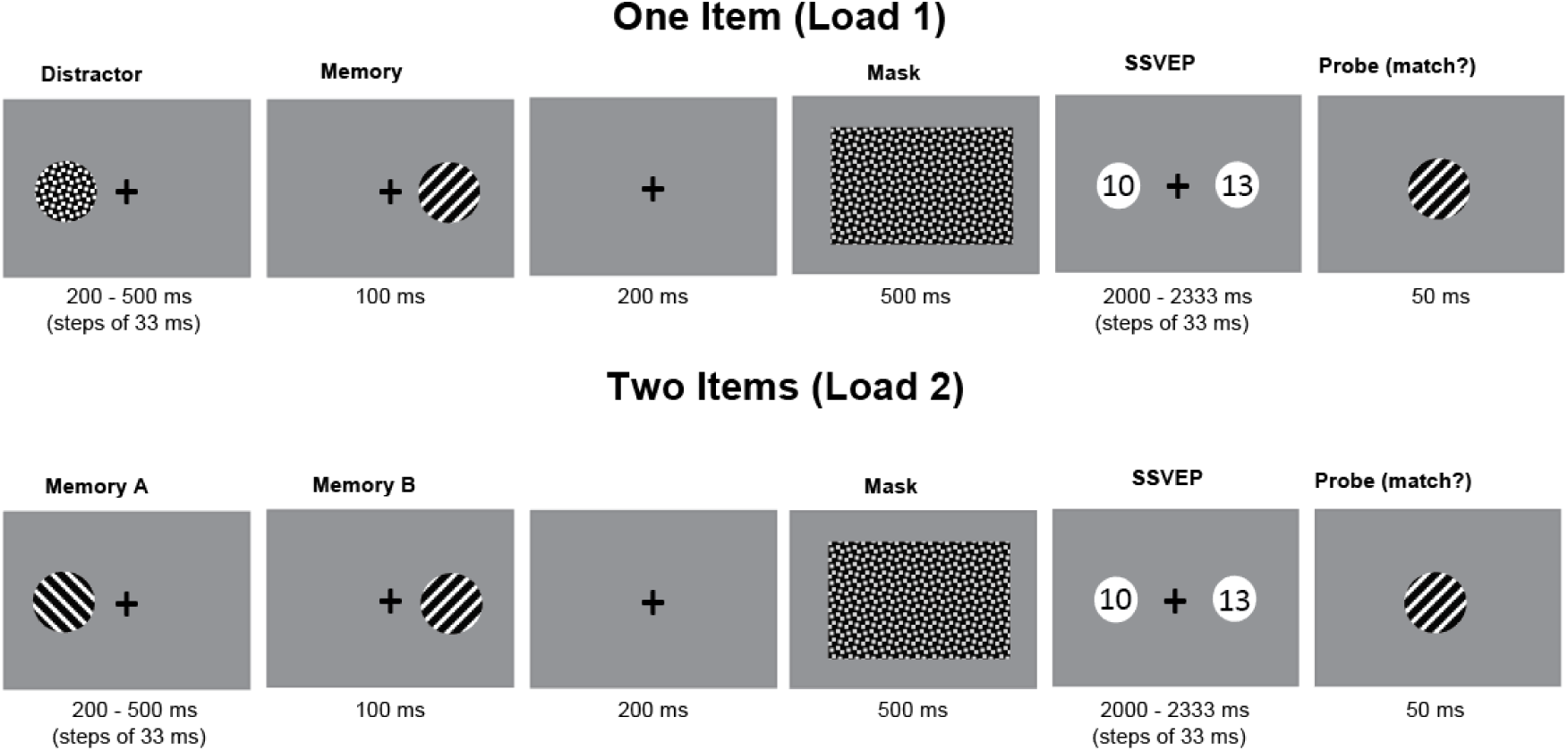
Participants performed a delayed match-to-sample task. In the one item (load 1) condition participants were presented with one to-be-memorized oriented grating and one distractor in random order and at a random location left or right of fixation. The two-item condition was identical to the one item condition, except that a second to-be-memorized memory item was presented instead of a distractor. Stimulus presentation was followed by a mask and subsequently two flickering discs were presented at the same location at which the initial stimuli were presented. Probes were presented at fixation and participants indicated whether the probes’ orientation matched either of the initial memory items.

To keep the task engaging and equally difficult between both conditions we used two online staircase procedures aimed at an average performance of 70%. This was done by adding gaussian noise to the probe stimulus depending on individuals psychometric functions estimated using Psychtoolbox QUEST algorithm (Farell & Pelli, 1999)

### Eye Tracking Recording and Analysis

Gaze position was continuously tracked using an Eyelink 1000 (SR Research, Ontario, Canada) eye tracker. A 13-point calibration was performed at the beginning of the experiment and after every 3 ^rd^ block. Gaze position was sampled at 1000 Hz.

Gaze data served to check for fixation throughout the memory and distractor presentation period (-1300 ms to -600 ms). Gaze position was baseline corrected by subtracting the median x and y coordinates during the baseline (-1800 ms to -1300 ms) from the entire trial. Trials were discarded from further analysis if fixation deviated more than 2° dva (x or y coordinates) from central fixation. Three participants were excluded from further analysis due to a large number of eye movements (>25% trials removed) during stimulus encoding. A re-analysis of the data including these 3 participants resulted in qualitatively identical findings.

### 1. 4. EEG recording and pre-processing

We recorded participants EEG using a 64 channel ActiveTwo Biosemi system. Two additional electrodes placed on the outer eye canthus and above the left eye recorded horizontal and vertical eye movements. Data analysis was performed in MATLAB using the Fieldtrip toolbox. Prior to all preprocessing steps we identified and removed bad channels via visual inspection. The EEG data were then re-referenced to the average of all channels, bandpass filtered between 0.5 and 80 Hz and line noise was removed using a DFT-filter (50 Hz). Thereafter the data were epoched from 2.3 s before SSVEP onset to 4 s after SSVEP onset. Large muscle related artifacts were first removed through visual inspection. Afterwards we performed an ICA on the datasets separately to remove eye-movement related artifacts. Finally, the data were down sampled to 512 Hz and absolute baseline correction was performed (window -1800 ms to -1300 ms before SSVEP onset).

### 1. 5. SSVEP and RESS

Frequency-specific SSSEP responses were isolated using a spatiotemporal source separation method (Cohen, 2022; Cohen & Gulbinaite, 2017), which is based on generalized eigenvalue decomposition (GED) and allows to maximize signal-to-noise ratio of steady-state responses by exploiting information present in inter-channel covariance matrices. Thus, instead of analyzing SSSEPs from a subset of electrodes with maximum power at the stimulation frequency, we analyzed a linearly weighted combination of signal from almost all electrodes. Notably, we removed a set (N=8) of frontal channels (FP1, FPz, FP2, AF7, AF3, AFz, AF4, AF8) before the creation of spatial filters as we expected a high amount of eye-movement related noise in these channels.

For each participant and stimulation frequency, a separate spatial filter was constructed by temporally narrow-bandpass filtering (Gaussian filter) the raw data (X) around the stimulation frequency f (10 Hz or 13.333 Hz, FWHM = 0.666 Hz) and at the two neighboring frequencies (f ± 0.666 Hz; FWHM = 0.666 Hz). Temporally filtered data (500 to 2000 ms relative to SSVEP onset) was then used to compute covariance matrices: one “signal” matrix (S covariance matrix) and two “reference” matrices that were averaged (R covariance matrix). The first 500 ms (0 to 500 ms) following SSVEP onset contain evoked potentials, and thus were excluded to not compromise the quality of the spatial filter (Cohen & Gulbinaite, 2017). Generalized eigenvalue decomposition (Matlab function eig) performed on “signal” and “reference” covariance matrices returned matrices of eigenvalues and eigenvectors. To increase the robustness of the spatial filters, we applied a 1% shrinkage regularization to the average “reference” covariance matrix. Shrinkage regularization involves adding a percent of the average eigenvalues onto the diagonals of the average “reference” covariance matrix (Cohen, 2022). This reduces the influence of noise on the resulting eigen decomposition. The eigenvectors (column vectors with values representing electrode weights, w) were used to obtain component time series (eigenvector multiplied by the original unfiltered single-trial time series, wT X).

The component with the highest signal-to-noise ratio in the power spectra at the stimulation frequency was selected for further analysis. The topographical representation of each component was obtained by left-multiplying the eigenvector by the signal covariance matrix (wTS). The obtained topographical maps were normalized and the sign of eigenvector was flipped for participants that showed spatial peaks opposite to that of the group average. The sign of the components affects only the representation of the topographical maps and has no effect on component time series (Cohen, 2022).

Differences in SSVEP responses associated with target and distractor locations were estimated by calculating Power and Coherence estimates at different stimulation frequencies. Power at each stimulation frequency was computed using FFT on trial-averaged component time series in the 500-2000 ms time window (relative to the SSVEP onset) and zero-padded to obtain frequency resolution of 0.066 Hz. The absolute value of FFT coefficients was squared and averaged across trials. To facilitate comparison across SSSEPs elicited by different stimulation frequencies, SSSEP power values were expressed in SNR units:

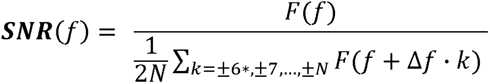

where N = ± 2 Hz, excluding 0.5 Hz around the frequency of interest. Individuals SNR values at 10 and 13.333 Hz were selected for further statistical assessment.

Time Frequency decompositions were performed via continuous wavelet transformation. The phase-locked power during SSVEP stimulation was calculated by multiplying the power spectrum of trial-averaged component time series with the power spectrum of complex Morlet wavelets 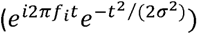, where *t* is time, *f*_i_ is frequency that ranged from 2 to 20 Hz in 0.333 Hz steps, and σ is the width of each frequency band defined as n/(2πf_i_), where n is a number of wavelet cycles which we set to n = 7.

Coherence was estimated between the single-trial component time series and pure sine waves (10 Hz and 13.333 Hz). First, we filtered 6300 ms epochs using a phase preserving, two-pass, Butterworth bandpass filters (4th order) with a hamming taper. The center filter frequencies were set to 10 Hz and 13.333 Hz respectively with a passband of ± 1 Hz. We then determined the analytic signal by the Hilbert transform which was subsequently used as the input for coherence at time point t (Eq. 2, adopted from Pan et al., 2021):

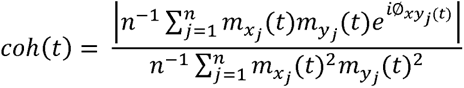

where *j* is the trial, *n* is the number of trials,*mx* (t) and *m_y_*(t) are the time-varying magnitude of the analytic signals from single-trial component time series and pure sine waves and ø*_xy_* (t) is the phase difference as a function of time between them.

### 1. 6. Statistical Analysis

To estimate the effect of internal attention on oscillatory power (SNR) in the one-item condition we first calculated attentional modulation indices (AMI, Zhigalov & Jensen, 2020) for both frequencies (10 Hz and 13.333 Hz, estimated with separate spatial filters) using the following formula:

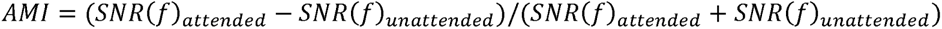

The resulting indices were then statistically tested using a two-way ANOVA with factors Load (1 and 2) and Frequency (10 Hz and 13.333 Hz) and t-tests.

Coherence time-series in one- and two item conditions were statistically assessed using a nonparametric cluster-based permutation test (Maris & Oostenveld, 2007). To this end we first calculated individual participants coherence time-series in the one item condition for memory and distractor locations (10 Hz and 13.333 Hz each) as well as both memory locations (10 Hz and 13.333 Hz) in the two item conditions. To test if coherence time-series significantly differed between (one/two-item) memory and one-item distractor conditions we first calculated the veridical difference in cluster-level t mass between two conditions. This was done by running t-tests for every individual timepoint of the group level difference time-series and identifying clusters of consecutive timepoints where the *p*-value fell below α = 0.05. The t-values within the largest cluster were summed to calculate the cluster-level t mass. We then randomly swapped 50% of labels in both conditions and recalculated the difference in largest cluster-level t mass. This procedure was repeated 10.000 times and generated the distribution that would be expected under the null hypothesis (H0: distribution of distance-to-bound score t-mass differences do not differ significantly between conditions). The veridically observed t-mass difference was compared to the null-distribution of t-mass differences and the null hypothesis was rejected if it exceeded the 95% quantile.

## Results

### 1. Load impacts behavioral performance

As expected, behavioral performance was lower with two to be memorized items than with one to be memorized item, expressing itself in a significantly lower hit rate (*t*(22) = 7.05, *p*<0.01), longer reaction times (*t*(22) = 5.52, *p* < 0.01) and higher false alarm rate (*t*(22) = 9.06, *p*<0.01) in the condition in which two items needed to be memorized.

**Figure 2.**
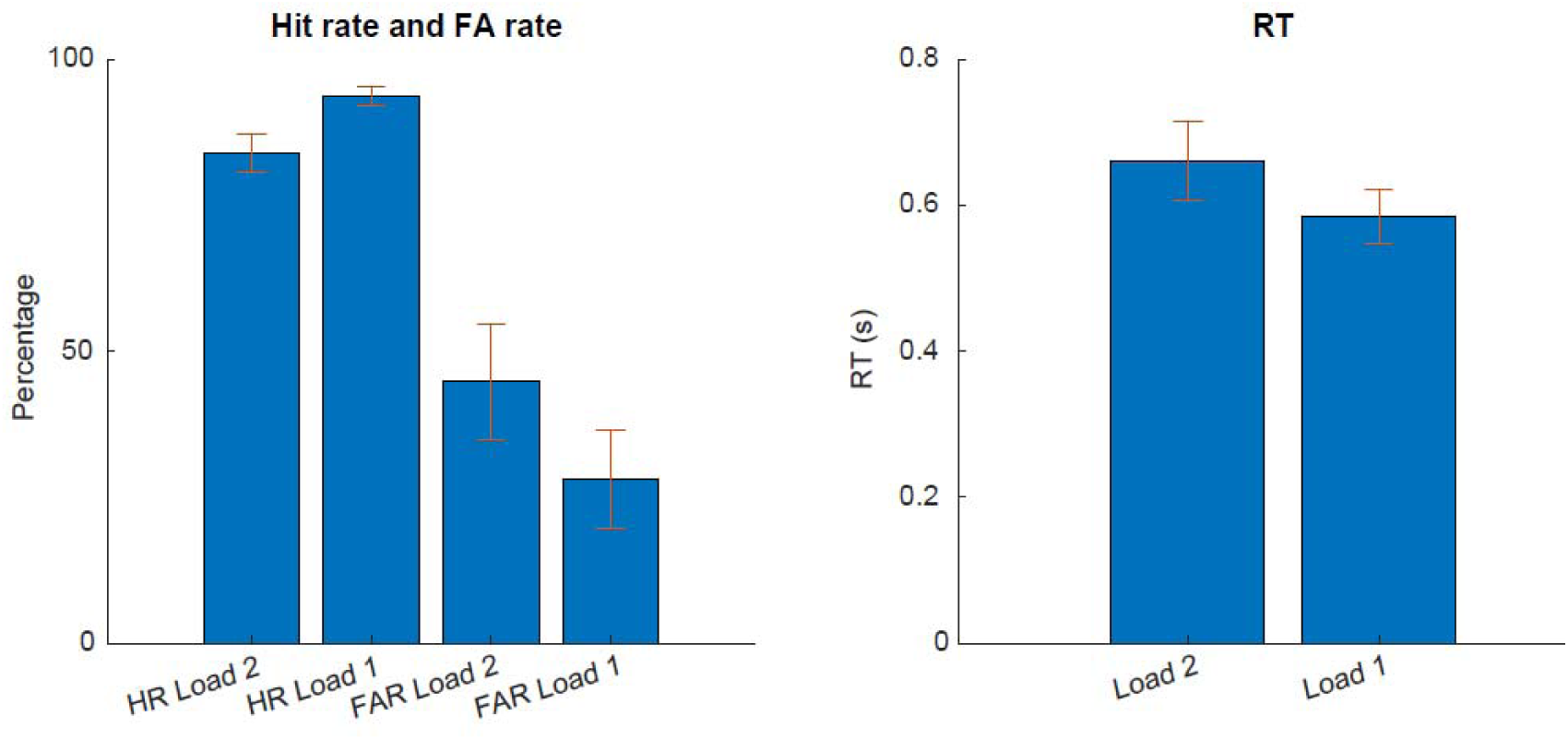
Behavioral Results. Group average performance (Hit rates and False Alarm rates) as well as reaction times for Load 1 and Load 2 conditions. Error bars indicate 95% confidence intervals.

### 1. 2. Dynamic allocation of spatial attention during VWM maintenance

We hypothesized that SSVEP responses corresponding to memory locations would be enhanced compared to distractor locations, reflecting the allocation of internal attention to the currently maintained VWM representation. Spectral analysis of 10 Hz and 13 Hz component time-series (500 ms to 2000 ms) revealed oscillatory peaks at the corresponding frequencies (Figure 4A,B).

Topographic representations of subject-average spatial filter maps for each SSVEP component (10 Hz and 13 Hz) show expected lateralization in scalp projections (Figure 4A,B).

To investigate the time-course of allocation of attention we compared SSVEP coherence time-series and time-resolved spectral power at frequencies corresponding to memory (Load 1 and Load 2) and distractor locations. Cluster-based permutation tests revealed significantly higher 13 Hz coherence at memory locations (Load 1, 1200 ms to 1950 ms; Load 2, 1350 ms to 2000 ms) compared to distractor locations (Figure 3B). We found no significant differences in 13 Hz coherence at memory locations between conditions Load 1 and Load 2. Furthermore, permutations tests revealed no significant differences between memory and distractor locations for 10 Hz coherence time-series (Figure 3A). To adjust for potential differences in individual’s absolute coherence we calculated normalized attentional modulation indices (AMI, see methods section 6. statistical analysis). Coherence values were averaged within significant time windows identified by the cluster-based permutation test 1350 ms to 1950 ms and AMI’s were calculated as follows: Load 1/Distractor: (Memory Load 1 – Distractor/ Memory Load 1 + Distractor), Load 2/Distractor: (Memory Load 2 – Distractor/ Memory Load 2 + Distractor) as well as the difference between the so calculated indices (Load 1/Distractor - Load 2/Distractor). As expected, attentional modulation indices (AMI) calculated from 13 Hz coherence was significantly larger than zero for Load 1/Distractor (*t*(22) = 3.97, *p* < 0.001) and Load 2/Distractor (*t*(22) = 3.01, *p* = 0.007) indicating a stronger allocation of attention to previous memory as compared to distractor locations. We found no significant differences in the AMI calculated from 13 coherence between Load 1 and Load 2 memory conditions (*t*(22) = 1.98,*p* = 0.06, Figure 3C) indicating that the degree of attentional allocation to a single location was not dependent on the total number of items in VWM. Furthermore, attentional modulation indices calculated from 10 Hz coherence did not differ significantly from zero in Load 1/Distractor (*p* = 0.72), Load 2/Distractor (p = 0.96) or when comparing between both (*p* = 0.75, Figure 3C).

**Figure 3.**
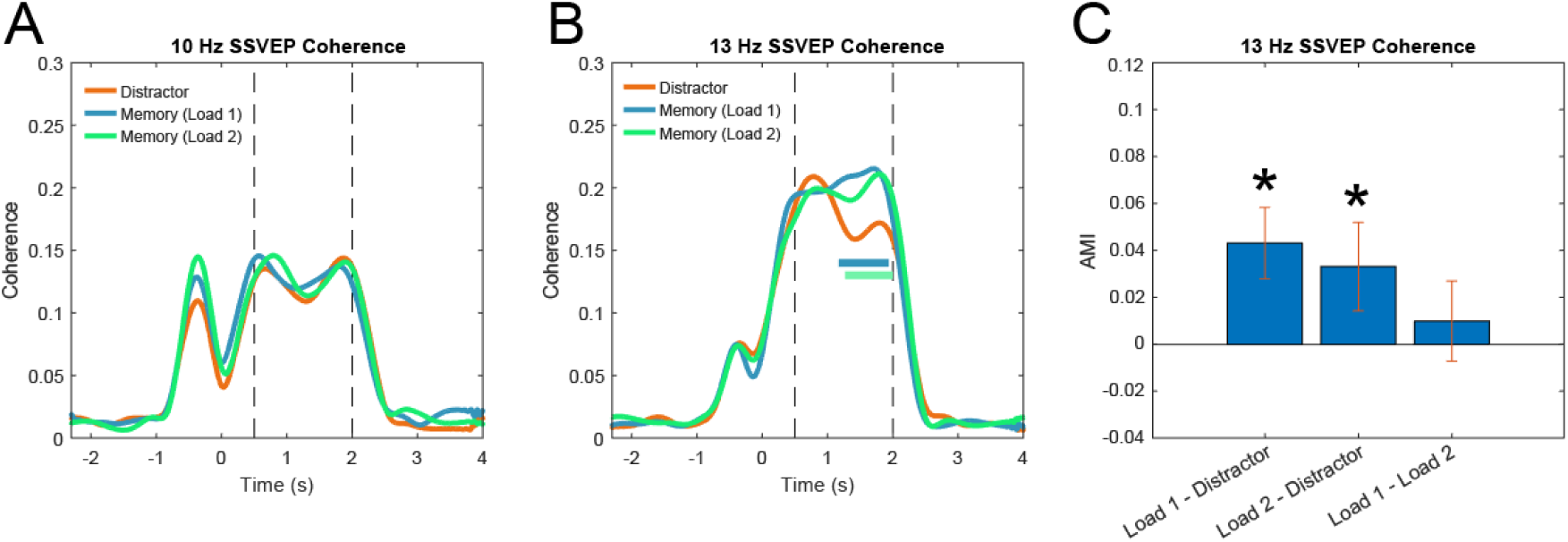
A. 10 Hz coherence over time. Orange line indicates coherence over trials in which the item at the 10 Hz tagged location was a distractor. Similarly, blue and green lines indicate coherence corresponding to 10 Hz tagged memory locations in Load 1 (bluea)nd Load 2 (green) conditions. Blue and green bars show clusters indicating significant differences between memory (load 1) and distractor locations (blue) and memory (load 2) and distractor locations (green). Black dotted lines indicate SSVEP time window of interest B. Same as A but for 13 Hz coherence. C. 13 Hz coherence AMI averaged within significant cluster (1350 ms to 1950 ms). Errorbars indicate 95% confidence interval. Coherence significantly drops for distractor against memory locations before probe presentation.

**Figure 4.**
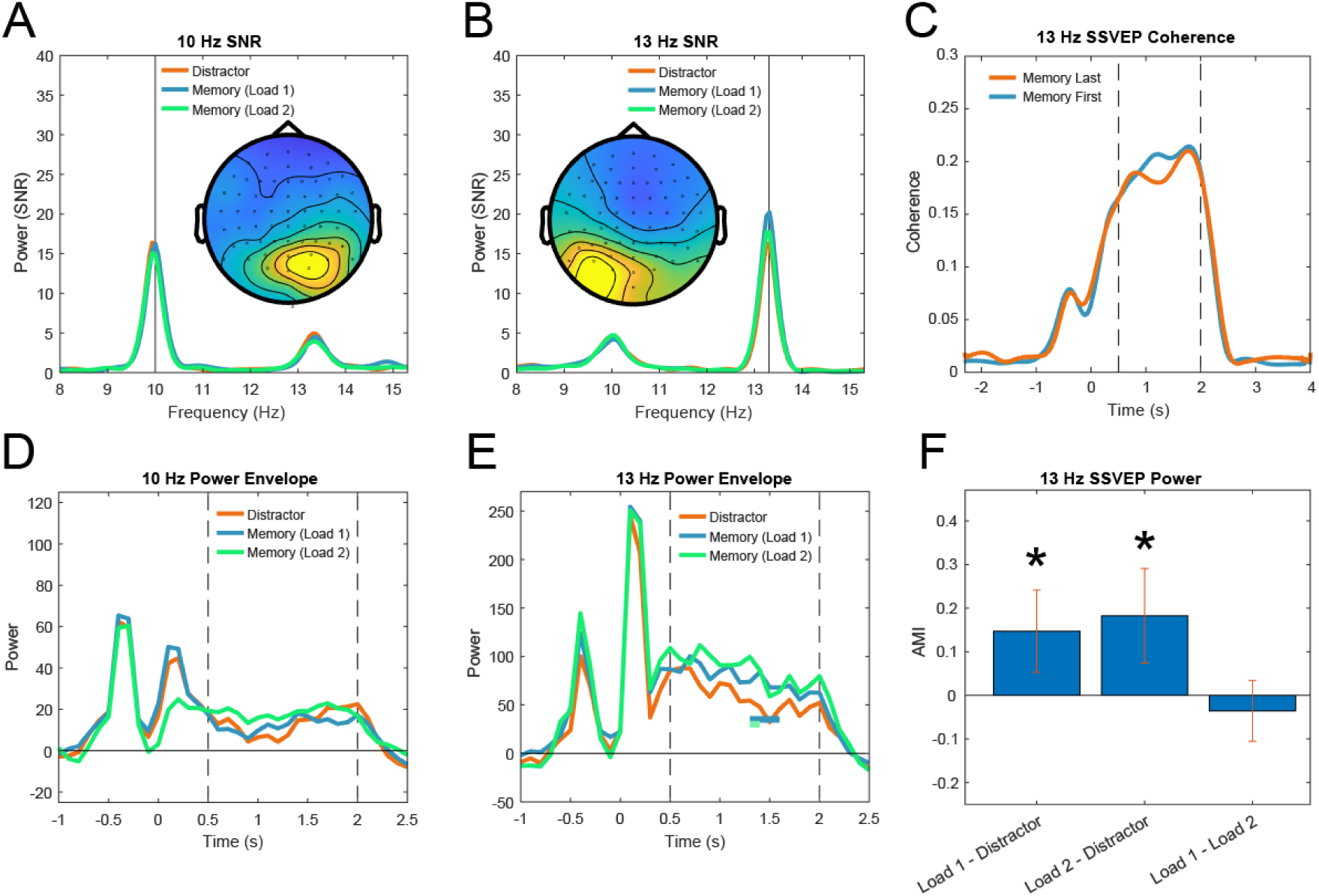
A,B. Signal to noise spectra of SSVEP component time series. Topographies display subject-averaged filter maps for 10 Hz and 13 Hz components respectively. C. 13 Hz coherence over time. Orange line indicates coherence over trials in which the item at the 13 Hz tagged location was presented last. Blue line indicates coherence over trials in which the item at the 13 Hz tagged location was presented first. D. 10 Hz power over time. Orange line indicates power in trials in which the item at the 10 Hz tagged location was a distractor. Similarly, blue and green lines indicate power corresponding to 10 Hz tagged memory locations in Load 1 (blue) and Load 2 (green) conditions. Blue and green bars show clusters indicating significant differences between memory (load 1) and distractor locations (blue) and memory (load 2) and distractor locations (green). Black dotted lines indicate SSVEP time window of interest E. Same as D but for 13 Hz power. F. 13 Hz power AMI averaged within significant cluster (1350 ms to 1950 ms). Errorbars indicate 95% confidence interval.

### Power

In addition to coherence, we analyzed the temporal dynamics of the SSVEP components by quantifying spectral power over time. This was done by extracting 10 Hz and 13 Hz power envelopes from corresponding time-frequency representations of component time-series (Figure 4D,E) and averaging power values centered around 10 Hz and 13.333 (±1 Hz) over time. Cluster-based permutation tests were performed to compare time-resolved power of memory (Load 1 and Load 2) and distractor locations. Permutations tests revealed significantly higher 13 Hz power at memory locations (Load 1, 1300 ms to 1500 ms; Load 2, 1300 ms to 1400 ms) compared to distractor locations (Figure 4E) but no differences in 10 Hz power between conditions (Figure 4D). Notably, the temporal profiles of 13 Hz coherence and power differed such that coherence generally increased over time whereas power decreased over time. Attentional modulation indices (AMI) calculated from 13 Hz power were significantly larger than zero for Load 1/Distractor (*t*(22) = 3.24, *p* = 0.004) and Load 2/Distractor (*t*(22) = 3.50, *p* = 0.002) but did not differ between the two (*t*(22) = -1.06*, p* = 0.30, Figure 4F). Furthermore, attentional modulation indices of 10 Hz components did not differ significantly from zero in Load 1 minus Distractor (*t*(22) = -1.38, *p* = 0.18), Load 2 minus Distractor (*t*(22) = 1.04, *p* = 0.17) or when comparing between both (*t*(22) = -1.75,*p* = 0.93, Figure 4F).

Taken together our findings show that 13 Hz SSVEP responses dynamically indexed the allocation of spatial attention to both memory and distractor locations. When spatial locations were previously occupied by a memory and a distractor item respectively, spatial attention remained on the memory location whereas it dropped off at the distractor location shortly before probe comparison. When two spatial locations were occupied by memory items, attention remained on both locations. Last, we found no differences in the amount of attentional allocated to individual locations when one versus two items were maintained in memory.

### 1. 3. Location specific effects on SSVEPs are not explained by attention during encoding

The effects of memory location on SSVEP responses could be explained either by sustained attention directed towards the item location in working memory or alternatively by persisting after-effects of spatial attention during stimulus encoding. We reasoned that potential after-effects of spatial attention during encoding should be stronger (or longer lasting) for locations where the last item was encoded as compared to locations where the first item was encoded. To test this prediction, we divided 13 Hz component time-series in the 2-item condition into two sets of trials. In the first set of trials memory items were encoded first at the 13 Hz stimulation location (memory first) whereas in the second set, memory items were encoded last at the 13 Hz location (memory last). Lingering after-effects of attention during encoding should hence manifest as stronger coherence in memory last trials. Cluster-based permutations tests revealed no significant differences between memory first and memory last, suggesting that location-specific effects on SSVEP coherence were not caused by potential after-effects of spatial attention during encoding.

### 1. 4. Reallocation of spatial attention predicts reaction times

The observed pattern of SSVEP responses indicates that spatial attention was maintained on both memory and distractor locations but reduced sharply at distractor locations shortly before the presentation of the probe in the center of the screen. We hypothesized that this reduction might be the result of a reallocation of attention from the distractor location to the center of the screen to facilitate perception of the probe. Simultaneously, attention might remain on the memory location as this could be a prerequisite for VWM maintenance.

To investigate whether the reallocation of attention from distractor to probe had a meaningful behavioral effect, we correlated individual participants coherence AMIs (Load 1/Distractor, Load 2/Distractor) at each timepoint with individual’s average reaction times. We hypothesized that a stronger reallocation of attention should lead to a more efficient perceptual processing of the probe stimulus and reduce reaction times. Coherence AMIs in the Load 1/Distractor and Load 2/Distractor conditions at multiple successive timepoints shortly before probe appearance (1700 ms to 1800 ms and 1600 ms to 1800 ms respectively) were significantly negatively correlated with reaction times (Figure 5B). These results suggest that attention was reallocated from the distractor location to the central probe location and that stronger shifts led to faster responses to the probe.

**Figure 5.**
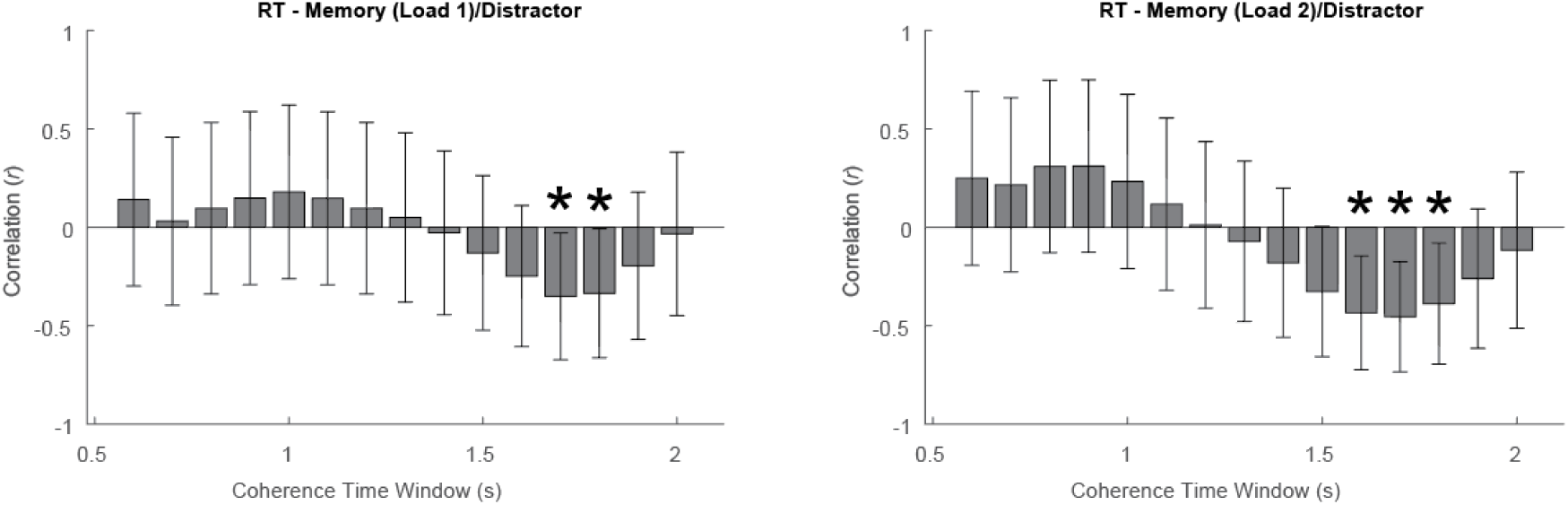
Pearson correlations between attentional modulation indices over time (100 ms bins) and average reaction times. Error bars indicate upper and lower 95% confidence intervals.

## Discussion

We here used SSVEPs to measure the dynamic allocation of attention to to-be-memorized gratings and to-be-ignored distractors memory items by exploiting that items were encoded (and maintained) spatially, even though space was irrelevant to answer the probe. We tested whether memory locations were attended stronger than distractor locations, how attention was distributed between more than one memory location and how attention was reallocated on a highly resolved timescale in anticipation of a central probe. Our findings show that SSVEP responses dynamically indexed the allocation of spatial attention to both memory and distractor locations. When both locations were occupied by memory items, attention was sustained on both. However, when one location was previously occupied by a distractor item spatial attention remained on the memory location but dropped off at the distractor location shortly before probe comparison. The degree of this reduction predicted reaction times – indicating that the reallocation of attention from irrelevant distractor to relevant upcoming probe location determines accurate retrieval and responding.

The central strength of our paradigm lies in its ability to demonstrate how attention is dynamically allocated to several relevant and irrelevant spatial locations during a visual working memory task. Recent work has highlighted the importance of studying working memory as a dynamic process that flexibly adjust to the current task goals (Buschman & Miller, 2022; Chota & Van der Stigchel, 2021; Nobre & van Ede, 2023). Previous behavioral work hereby relied primarily on probe stimuli to measure item-location specific attentional effects (Awh et al., 1998; Awh & Jonides, 2001; Chen & Wyble, 2015; Elsley & Parmentier, 2015; Theeuwes et al., 2011). While informative about the effects on perception, this approach is inherently limited in its ability to uncover dynamical changes in attention as only single timepoints can be probed per trial. Experiments using classic electrophysiological measures such as alpha lateralization (Ede et al., 2017; Foster et al., 2017; Liu et al., 2022, 2023; Poch et al., 2014, 2017) or time-resolved behavioral measure such as eye-movements (van Ede et al., 2019, 2020, 2021) attempt to paint a more dynamic picture of attention but ultimately only provide correlational evidence for its effects on perceptual processing. In contrast, the here utilized SSVEP paradigm provides a highly time-resolved and functionally direct estimate of how attention modulates spatially specific perceptual processing. Moreover, we could measure attentional allocation without using behavioral probes at memory or distractor locations. This uniquely allowed us to test attentional facilitation without participants explicitly directing spatial attention to potential probe locations and hence isolate memory-specific attention.

One of the more surprising results in this experiment is the fact that distractor locations were attended for almost 2100 ms after the distractor was presented. We propose several possible explanations for this which might not be mutually exclusive. First, participants might have accidentally encoded distractors in VWM due to attention automatically being captured by the abrupt onset, the relatively short presentation time (300 to 600 ms) and no prior knowledge on the upcoming location of the relevant memory item. Previous work showed such occasional encoding and maintenance of distractors for more than a second, even with predictable spatial location (Vogel et al., 2005). The sustained increase in perceptual processing at distractor locations might therefore index a similar engagement of spatial attention as found during memory maintenance. While maintaining distractors attended in VWM might seem counterintuitive, the active removal of items from VWM and hence from the focus of spatial attention could be more effortful than simply maintaining them in memory. Only when attentional resources are required elsewhere, in our case when probe presentation is imminent, attention would then be reallocated from the encoded distractor to more relevant locations. An additional explanation for the sustained attention directed to distractor locations is that the flickering stimuli themselves exogenously attracted attention which led to a saturation of the SSVEP response measured on the scalp. This could also explain why we do not see a general reduction in SSVEP coherence in the two item condition in which attentional resources also would have to be dedicated to the probe location shortly before its appearance. Under this assumption the reduction in SSVEP coherence at the distractor location might reflect the active suppression or removal of the accidentally encoded distractor item from memory, leading to a strong reduction in attention directed to its former location. Similar to our initial hypothesis this would also provide a parsimonious explanation for the link between SSVEP responses and reaction times, as the inhibited item might free up working memory capacity for the upcoming probe. In their essence these two explanations point to two distinct models of attention in working memory: In the first, attentional resources are deliberately allocated and reallocated between encoded items. In the second, available attentional resources are automatically distributed between all encoded items and reallocation entails the active inhibition and/or removal of items from the pool of memories. Future work, will have to dissociate between these two models by adding flicker locations at which neither memory nor distractor items are presented.

To our knowledge there is only a single study that has used SSVEPs to probe memory and distractor locations during the maintenance interval (Vissers et al., 2017). The authors found no attentional effect of memory versus distractor locations at the entrained frequencies of 16/18 Hz but revealed larger amplitudes for memory locations at the second harmonic (32/36 Hz), in line with our results. They hypothesized that the attentional effect might not have been revealed at the entrainment frequencies because the placeholders instead of precise stimulus locations were flickered. This might have led to a surround suppression and subsequently a reduction in amplitude of the flicker signal. Notably the probes in their study were presented at locations that overlapped with the memory/distractor items – spatial attention to the memorized items and to the upcoming probes can therefore hardly be dissociated. Although Visser and colleagues (2017) did not statistically assess dynamic changes in attention during the maintenance period, visual inspection of the SSVEP signal over time indicates a reduction in attention over distractor locations ∼1000 ms before probe onset which tightly resembles the attentional reallocation demonstrated in our work. Together with the presently reported data, this provides converging evidence that spatial attention dynamically modulates SSVEP responses during VWM maintenance.

How are attentional resources distributed between one or more relevant memory locations during maintenance? Previous work has shown that SSVEP amplitude is reduced when attention is distributed between two instead of one relevant location (Andersen et al., 2009; Toffanin et al., 2009). In contrast, we found no differences in SSVEP responses between one-item and two-item conditions. This discrepancy might be due to both studies using external distractors that had to actively be ignored. When presented with two competing streams of external information the visual system might benefit from dedicating additional resources to the attended location. Spatial attention might therefore be redistributed only to the attended visual stream leading to increased SSVEP responses. In our study there was no competing stream of external visual information that had to be actively suppressed, which might have made it less important to redirect additional attentional resources to the relevant memory location. This interpretation is in line with the hypothesis that attention plays an important role in protecting memorized and perceived neural representations from external interference (Souza & Oberauer, 2016). Future studies might introduce more interfering distractors in the maintenance period to test if attentional allocation changes as a function of external distractor load.

It could be brought forward that the location-specific effects observed here are not specific to memory maintenance or retrieval but rather reflect lingering attentional effects caused by stimulus encoding. We believe this to be unlikely: First, one would expect encoding related effects to be most prominent at the location of the last item and weakest at the location of the first item. We explicitly tested this by comparing coherence between trials in which the first or the second item was presented at the 13 Hz SSVEP location finding no effect of encoding order. Second, attentional facilitation caused by attentional orienting to peripheral cues has been shown to peak around 175 ms and does generally not last longer than 400 ms (Müller & Rabbitt, 1989) whereas we found the strongest facilitation to emerge at around 1800 ms.

While we observed a consistent effect of memory/distractor location on 13 Hz SSVEP responses we did not find similar effects for 10 Hz responses. This might be explained by the fact that 10 Hz entrainment interferes with the brain’s endogenous alpha rhythms which can be found at similar frequencies (Gulbinaite et al., 2017, 2019). Previous work has shown that either an increase or decrease in flicker amplitude in the alpha range can be observed depending on the individual peak alpha frequency of the participant and the strength of endogenous alpha oscillations (Ding et al., 2006; Gulbinaite et al., 2019; Wang et al., 2007). Endogenous alpha oscillations have been demonstrated to decrease over visual areas were memory items were encoded (Liu et al., 2023; Poch et al., 2014, 2017). The lack of an effect at 10 Hz might therefore be the result of increasing responses to the flicker stimulus and decreasing endogenous alpha oscillations cancelling each other out.

Our results provide further evidence to the idea that visual memoranda are stored in a spatially organized format (Schneegans & Bays, 2017; van Ede et al., 2019). Furthermore our findings are in line with the sensory recruitment hypothesis claiming that visual sensory areas, which are largely retinotopically organised, are involved in the maintenance of memoranda (Ester et al., 2009, 2016; Harrison & Tong, 2009; Rademaker et al., 2019; Serences et al., 2009). The spatial organization of the visuo-attentional system is assumed to play a key role in the perceptual binding of multiple features into objects (Kahneman et al., 1992; Treisman, 1988). Our findings support the idea that this also extends to objects stored in memory because spatial attentional effects could be observed despite “external” memory locations remaining completely irrelevant throughout our experiment.

In conclusion, we show that SSVEPs are a powerful new tool to study dynamic changes in internal attention directed towards individual representations in VWM. This was only possible as spatial locations of encoded items were irrelevant to answer probes – supporting the notion of a spatially organized VWM. We demonstrate that VWM maintenance and utilization is accompanied by dynamic, location-specific and behaviorally relevant (re-)distribution of attention indexed by SSVEP coherence. Crucially, while spatial attention is continuously maintained on memory locations it is reallocated from distractor to a central location to facilitate encoding of the probe leading to faster reaction times.

## Notes

### Competing Interest Statement

The authors have declared no competing interest.

